# Additive manufacturing of patient-specific intracranial aneurysm cell culture models

**DOI:** 10.1101/2025.02.24.639980

**Authors:** Chloe M. de Nys, Ashley R. Murphy, Jaimee-Lee Wood, James I. Novak, Danilo Carluccio, Craig D. Winter, Mark C. Allenby

## Abstract

Intracranial aneurysms (IAs) are present in 2-6% of the global population. While rare, rupture results in mortality rates of 30-50% and lifelong disabilities in survivors. While treatment of unruptured IAs carries its own risk of mortality, there are no rigid guidelines indicating which IA presentation is at greater risk of rupture. We develop and evaluate the suitability of various additive manufacturing processes to fabricate patient-specific IA culture models for understanding IA pathophysiology and thereby support future development of a rupture risk prediction tool.

Material compatibility of several 3D printed resins, polydimethylsiloxane (PDMS) and collagen gel with immortalised human brain endothelial cells (HBECs) were investigated. Patient angiograms were segmented to produce *in vitro* models via two fabrication approaches: stereolithography (SLA) 3D printing versus chocolate injection moulding of a sacrificial core embedded in PDMS. These 3D arterial models were then cellularised with HBECs, and geometric accuracy and distension properties evaluated.

PDMS, collagen gel and the elastic50A resin materials supported cellularisation on material surfaces with high cell viability and proliferation. Both the 3D printed resin and injection moulding techniques successfully fabricated patient-specific basilar artery IA models with a Dice-Sørensen coefficient of over 90%. However, only PDMS models offered complete cell coverage in 3D geometries. This exceeds current benchmarks on IA fabrication accuracy. Through controlling IA model wall thickness, we demonstrate the ability to create localised distension under pressure in the context of thin– and thick-walled aneurysms.

*In vitro* IA models present a promising platform for investigating IA pathophysiology and rupture risk. The novel chocolate sacrificial core technique offers a biocompatible, support-free fabrication method suitable for 3D cultures. Given the independent effects of fluid dynamics and mechanical strain on cell behaviour, it is essential to characterise distension under pressure and ensure accurate fabrication for reliable analysis into cell behaviour.

## 1. Introduction

Intracranial aneurysms (IAs) are an outpouching of the intracranial arterial wall, with an estimated global prevalence of 2-6% [1–4]. Up to 1.1% of IAs will rupture to cause a subarachnoid haemorrhage, resulting in a 30-50% mortality rate with 50% of survivors left with permanent disabilities [1, 2, 5, 6]. If detected early, at-risk IAs can be treated through surgical clipping or endovascular coiling. Both treatment options pose their own procedural risks, with treatment related fatality and morbidity of up to 5% [7, 8]. As such, clinicians must evaluate and assess the risk of a rupture event against procedural risks to determine the most appropriate patient treatment. However, no universal definitive IA rupture risk prediction tool exists. Clinicians are currently reliant on simple risk prediction methods that consider morphological and patient characteristics, and their clinical experience to determine the necessity for surgical treatment.

The most common risk prediction method currently employed is termed PHASES, which considers the patient Population, Hypertension, Age, Size of Aneurysm, any Earlier subarachnoid haemorrhage and the Site of the aneurysm [9]. While other proposed risk prediction methods such as the Unruptured Intracranial Aneurysm Treatment Score (UIATS) considers additional factors such as aneurysm morphology and lifestyle habits, it has proved unreliable in providing clear recommendations for treatment in patients whose aneurysms eventually ruptured [9, 10]. However, both scoring systems have limited universal value given the lack of correlation with clinical opinion and professional preference [9, 11, 12].

Where there remains no comprehensive method for discriminating between stable IAs and those prone to rupture, further research into IA pathophysiology is required. Limited knowledge in human IA pathophysiology to-date could be attributed to the rarity of clinical IA samples obtained either post-mortem or intra-operatively, with most studies evaluating IA stability relying on imaging technologies [13].

As such, *in vivo* animal models have been heavily utilised in IA research since the first canine model in 1961 and the more commonly used rabbit-elastase model of 1963 [14, 15]. These models require surgical manipulation or elastase to degrade and weaken intracranial arteries [15] and therefore may not accurately allow translation to human IA growth and rupture mechanisms [16]. Additionally, many risk factors influencing human IA development and rupture such as smoking and hypertension, cannot be easily incorporated into these models.

Alternatively, novel *in vitro* bioengineered models could provide a means to study IA remodelling and rupture mechanisms and evaluate new clinical therapies. One of the first endothelialised *in vitro* IA models was developed by Kaneko et al. (2018) utilising a sacrificial core made of acrylonitrile butadiene styrene (ABS) adopting a patient specific geometry [17]. This was coated in polydimethylsiloxane (PDMS) and then dissolved, resulting in a hollow model. Similarly, Levitt et al (2019) fabricated a planar PDMS aneurysm model around a stereolithography (SLA) printed resin core that was mechanically removed [18]. These *in vitro* IA models were then cellularised and perfused, confirming changes in cell morphology, alignment, and gene expression between the parent vessel and IA domes, which also correlated with fluid flow dynamics [17]. More recently, Yong et al (2021) succeeded in creating an idealised IA model in gelatine methacrylate (GelMA), co-cultured with human primary brain vascular smooth muscle cells and human umbilical vein endothelial cells [16]. Long-term perfusion culture of 10 days was achieved with both cell types maintaining their secretome profiles based on analysis of conditioned medium [16]. These studies demonstrate the potential of *in vitro* bioengineered models as a novel avenue to analyse cellular remodelling in IAs, with the ability to correlate cell behaviour with IA morphological predictors and blood flow mechanics.

However, variations in existing fabrication techniques complicates direct results comparisons, as model design and material properties can influence cell behaviour, haemodynamic parameters, and cyclic stretching. For example, the geometric accuracy of existing *in vitro* IA models is unknown, where small variations can significantly affect biofluid flow dynamics [19]. Similarly, model compliance can also alter flow profiles, where a rigid model can significantly increase peak wall shear stress [20]. Material stiffness is well known to influence cell proliferation and phenotype [21], while mechanical stretching alone can influence cellular inflammation regardless of haemodynamic stress [22]. Furthermore, while 3D printed IA models are frequently created for surgical training and endovascular simulations [23–26], and several 3D printed materials have been evaluated for biocompatibility [27], the use of 3D printed resin materials to fabricate IA models for cell culture has not yet been demonstrated.

Therefore, this work aims to evaluate the suitability of various additive manufacturing processes to create *in vitro* IA models that can support the viable culture of phenotypical human brain endothelial cell (HBECs) monolayers. Material biocompatibility, geometric accuracy and distension properties of patient-specific IA models will be characterised to support the implementation of *in vitro* platforms into future research investigating IA pathophysiology.

## 2. Methods

### 2.1 Material fabrication and post-processing

Elastic50A (V1, Cat. No: RS-F2-ELCL-01) and Flexible80A (Cat. No: RS-F2-FL80-01) resins were printed using the Form 3/3+ stereolithography printer (Formlabs, 0.1 mm z-resolution) and post-processed as per the supplier’s instructions. Briefly, the printed resins were washed in a clean beaker of fresh 100% isopropyl alcohol (IPA) for 10 minutes, then transferred to a new beaker of fresh 100% IPA for a further 10 minutes. Printed resins were then placed into the Formlabs Form Cure and exposed to 405 nm light at 60°C for 20 minutes and 10 minutes for Elastic50A and Flexible80A respectively. The newly released BioMed Elastic50A resin (Cat. No: RS-C2-BMEL-01) was also printed on the Form 3/3+ printer and post-processed as per the supplier’s instructions, for use in preliminary 2D and 3D culture studies.

Agilus30^TM^ and ‘Compliant Blood Vessel’ (combination of VeroClear^TM^ and Agilus30^TM^) resins (Stratasys) were printed on the Stratasys J750 Digital Anatomy PolyJet printer (14 – 27 µm z-resolution) with supports printed in SUP706 (Stratasys). Supports were manually removed before soaking in a caustic solution (2% sodium hydroxide, 1% sodium metasilicate) using the CSIIP CleanStation (Stratasys) for two hours to remove remaining supports.

Sylgard^TM^ 184 Silicone Elastomer kit (Polydimethylsiloxane or PDMS, Cat. No: 500GM184KIT) was used to prepare PDMS by mixing the Elastomer base with the curing agent in a 10:1 ratio and cured at room temperature for 24 hours. PureCol^®^ Type I Bovine Collagen Solution (3 mg/mL, Cat. No: 5005, Advanced Biomatrix) was prepared as per the supplier protocol for 3D gel preparation at a working concentration of 2.4 mg/mL.

### 2.2 Maintenance culture of HBEC-5i (HBECs)

Human cell culture was supported by The University of Queensland Human Research Ethics Committee 2021/HE002698 and Institutional Biosafety Committee approval IBC/602E/ChemEng/2023. Human cerebral microvascular endothelial cells (HBEC-5i, CRL-3245, ATCC^®^, USA) were cultured in media comprising Dulbecco’s Modified Eagle Medium/Nutrient Mixture F-12 (DMEM/F12, Cat no: 11320033) supplemented with 10% (v/v) Fetal Bovine Serum (FBS, Cat. no: 10099141) and 1% endothelial cell growth supplement (Cat. No: E2759, Sigma Aldrich, USA). 1% (v/v) penicillin-streptomycin (PenStrep, Cat. No: 15140122) was added when used in 2D and 3D material cultures. Cells were cultured at 37°C in a humidified atmosphere of 5% CO_2_ in air below passage 10 for all experiments.

### 2.3 Seeding HBECs on 2D materials

Each resin was 3D printed into 1 mm thick rectangular slabs directly on the print beds with no angulation. PDMS samples were prepared by pouring the mixture into a clean petri dish with a 1 mm thickness before curing at room temperature for 24 hours. Resin and PDMS disks were created using a 5 mm biopsy punch which were then glued to 6 mm glass circular microscope coverslips with Sika Aquarium SikaSeal glue (Cat. No: 1210603, Bunnings, Australia). Glued samples were left at room temperature for seven days to cure. 200 µL of collagen solution was deposited directly into wells of a 48-well plate before curing at 37°C for two hours to facilitate complete gelation.

To prepare materials for cell seeding, resin and PDMS samples were rinsed with 70% (w/v) ethanol then distilled water and allowed to dry before oxygen plasma treatment at 40 W for 60 seconds. Samples were then placed into wells of a 48-well plate under aseptic conditions. Samples were sterilised by soaking in 70% (w/v) ethanol for 10 minutes and exposed to germicidal ultraviolet light for 30 minutes. Resin, PDMS and collagen gel samples were then washed twice with phosphate buffered saline (DPBS, Cat. No: 14190144) for 5 mins and incubated at 37°C with 0.1% gelatine (Cat. No: 07903, Stem Cell Technologies, Australia) overnight. Gelatine was then aspirated, and HBECs seeded at 20,000 cells/cm^2^. Media exchanges were performed every two days for up to 14 days total.

### 2.4 Material cell viability

Cell viability was assessed at day two and day four of culture using a Live/Dead Cell Double Staining Kit (Cat. No.: 04511, Sigma-Aldrich). A staining solution of 2 µL of Calcein-AM, 1 µL propidium iodide and two drops of NucBlue^TM^ Live ReadyProbes^TM^ Reagent (Hoechst 33342, Cat. No: R37605) was prepared in 1 mL of DPBS to stain live cells, dead cells, and the cell nuclei respectively. Samples were incubated with this staining solution for 15 minutes at 37°C before imaging using an Olympus IX83 fluorescence microscope. Images were processed using a linear brightness and contrast adjustment, and analysed using ImageJ [28].

### 2.5 Immunofluorescent staining

Samples were fixed in 4% paraformaldehyde (Cat. No: C006, ProSciTech Australia) for 10 minutes at day 7 and day 14. Due to autofluorescence of the elastic50A resin under ultraviolet light (405 nm), 0.1% Sudan Black B (SBB, Cat. No: 199664, Sigma-Alrich) prepared in 70% (w/v) was applied to elastic50A samples for three hours at room temperature. Samples were washed with 70% (w/v) ethanol until SBB residue was no longer present, before rinsing with DPBS. All samples were then permeabilised with 0.1% (v/v) Triton^TM^ X-100 solution (Cat. No: 93443, Sigma-Aldrich) for 10 minutes, washed with DPBS and blocked using 10% (v/v) goat serum (Cat. No: 16210064, Thermo Fisher Scientific) in DPBS for 60 minutes at room temperature.

Immunofluorescent (IF) staining was performed in the dark using anti-Ki67 Rat IgG2a primary antibody (5 µg/mL, SolA15, eBioscience^TM^, Cat. No: 14-5698-82) and the Rat IgG2a kappa Isotype Control (eBR2a, eBioscience^TM^, Cat. No: 14-4321-82) at room temperature for three hours, followed by goat anti-Rat IgG Alexa Fluor^TM^ 555 secondary antibody (4 µg/mL, Alexa Fluor 555, Cat. No: A-21434) for one hour at room temperature. Samples were then stained with NucBlue^TM^ Fixed Cell ReadyProbes^TM^ Reagent (2 drops per mL in DPBS, Hoechst 33342, Cat. No: R37605) and ActinGreen^TM^ 488 ReadyProbes^TM^ Reagent (2 drops per mL in DPBS, Cat. No: R37110) at room temperature for 30 and 60 minutes, respectively.

Fluorescence images were acquired using a Nikon Eclipse Ti2 fluorescent microscope and a linear brightness and contrast adjustment was applied. Exposure time was doubled for elastic50A samples under the FITC and TRITC channels to account for residual autofluorescence and reduced signal intensity due to the SBB. Images of elastic50A also required background subtraction in addition to the linear brightness and contrast adjustment (refer to Supplementary figures S1 and S2) to ensure visualisation and quantification using ImageJ [28].

### 2.6 Patient cranial scans with intracranial aneurysms

De-identified medical images of patients with IAs acquired through Metro North Hospital and Health Service (MNHHS) between 2008 and 2022 were collected for research purposes as approved by the Royal Brisbane and Women’s Hospital (RBWH) Human Research Ethics Committee (Ref: LNR/2019/QRBW/49363) and the Queensland Department of Public Health (Ref: Public Health Agreement #49363). Data acquired includes digital subtraction angiography (DSA), computed tomography angiography (CTA) and time-of-flight magnetic resonance angiography (TOF-MRA) images. The TOF-MRA dataset has previously been published to the open-access repository OpenNeuro (doi:10.18112/openneuro.ds005096.v1.0.3) [29].

From this dataset, sub-051 (filename: sub-051/ses-20210702/anat/sub-051_ses-20210702_acq-tra-angio.nii.gz) was selected for modelling of a middle cerebral aneurysm (MCA) (4 × 2 × 3 mm) using an MRA scan. A DSA scan of a patient with a large basilar aneurysm (BA) (12 × 11 × 12 mm) not yet published in this dataset was also selected for modelling. Asymptomatic brain vasculature of patients IXI-017-Guys-0698-MRA and IXI019-Guys-0702-MRA from the online IXI (http://brain-development.org) database was used for segmentation and modelling of the basilar artery and middle cerebral artery respectively. Scans were downloaded in their NIFTI (Neuroimaging Informatics Technology Initiative) format and converted to DICOM (Digital Imaging and Communications in Medicine) format using an online NIFTI to DICOM converter (https://www.onlineconverter.com/nifti-to-dicom).

### 2.7 Segmentation and design of aneurysm and arterial models

DICOMs were imported into Mimics software (Materialise NV, v25.0) and segmented by pixel thresholding for the three-dimensional reconstruction of the lumen of vessels throughout the cranial vascular tree. The arterial segments of interest were then isolated. The smaller pontine arteries branching from the basilar artery were removed due to their modelling complexity and reduced resolution. Arterial segments were then smoothed and wrapped before exportation to a stereolithography (STL) model (Figure 1-Segmented). Using 3-matic software (Materialise, v25.0), artificial extensions were added to the ends of the arterial segments to support the attachment of barbed connectors and plugs upon fabrication (Figure 1-Extended).

**Figure 1:**
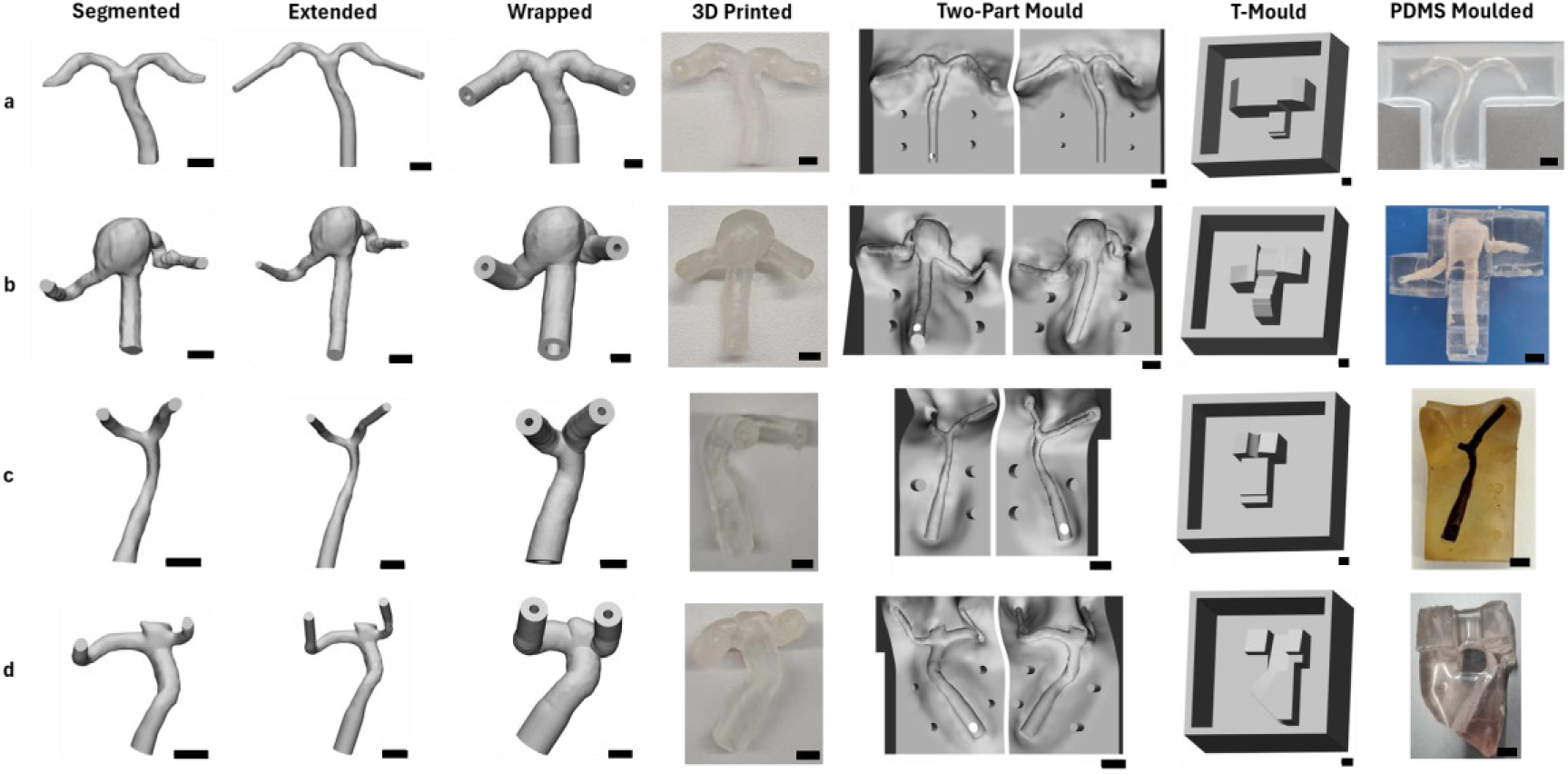
Modelling and fabrication of intracranial arterial and aneurysm models. (a) Asymptomatic basilar artery. (b) Basilar artery aneurysm. (c) Asymptomatic middle cerebral artery. (d) Middle cerebral artery aneurysm. A PDMS moulded model of the asymptomatic middle cerebral artery could not be created due to the bifurcation being prone to breakage as pictured. Scale bars represent 5 mm

3D *in vitro* aneurysm and arterial models were fabricated by either direct 3D printing, or by injection moulding a sacrificial core and embedding this into a surrounding biocompatible material. To create a hollow model for direct 3D printing, the arterial segment was ‘wrapped’ with a thickness of 1– and 2-mm. Then, the arterial segment was subtracted from the ‘wrapped’ geometry, and the ends of the model cut to yield a hollow arterial geometry with a 1-or 2-mm wall thickness (Figure 1-Wrapped).

To create an arterial sacrificial core (e.g. meltable or dissolvable) a two-part injection mould (Figure 1 – Two-Part Mould) was created. This was done by manually moulding a parting line around arterial segments using the ‘push/pull’ function in 3-matic software. A funnel was added to allow for filling of a sacrificial material through one side of the mould. For embedding the solid sacrificial arterial core into a surrounding biocompatible material, a separate T-shaped mould was designed around the arterial segment (Figure 1-T-Mould) and 3D printed in PLA using the Ultimaker S3 Fused Deposition Modelling printer. Designed as a positive mould, this was filled with PDMS to create a negative mould that could house the sacrificial arterial core with the correct orientation. These PDMS T-moulds were then lightly sprayed with mould-release spray (Ease Release 200, Art Tree Creations) to prevent the adhesion of filled PDMS.

### 2.8 Aneurysm and arterial model fabrication

#### 2.8.1 Direct 3D printing

Hollow arterial models (Figure 1-Wrapped) were directly printed without internal supports in the Elastic50A and BioMed Elastic50A resin using the Form 3/3+ stereolithography printer (0.1 mm z-resolution). External supports were mechanically removed before post-processing, as described in section 2.1.

#### 2.8.2 Injection moulding of a sacrificial core

Two-part injection moulds (Figure 1-Two-Part Mould) were printed in the Flexible80A resin using the Form 3/3+ stereolithography printer (0.05 mm z-resolution). External supports were mechanically removed before post-processing as described in section 2.1.

Injection moulds were sprayed with mould release spray before use. Sacrificial cores were created by injecting tempered cooking chocolate (Nestle Plaistowe Dark Cooking Chocolate, 40% Cocoa) into the two-part moulds, and left to set at room temperature. The moulds were carefully separated, and arterial cores removed. Any defects, such as the parting line, were carefully corrected using a warmed spatula to smooth the surface. Chocolate arterial cores were then kept at 4°C before being placed into the negative PDMS T-mould and surrounded with PDMS. Moulds were left to cure at room temperature for 24 hours.

Once cured, PDMS models inside the T-moulds were carefully removed. The sacrificial chocolate arterial cores were washed out using warmed water with the aid of sonication (Figure 1– PDMS Moulded). A chocolate core could not be created for the asymptomatic middle cerebral artery model due to the small diameter at the bifurcation being prone to breakage. An example of the broken chocolate artery in the two-part mould is presented in Fig 1c – PDMS Moulded.

### 2.9 Analysis of geometric accuracy and distension

Fabricated models were imaged using the Yxlon FF35 CT system to evaluate geometric accuracy and distension behaviour. Models were exposed to a direct transmission beam at 78.87 kV, 281.51 µA and 22.11 W with an exposure time of 181.82 ms, producing a voxel pitch of 33.33 µm. Scans took approximately eight minutes to complete.

The distension behaviour of BA aneurysm models fabricated in PDMS (Fig 1b – PDMS Moulded) and Elastic50A resin (Fig 1b – 3D Printed) with a 1 mm and 2 mm wall thickness were compared. Each model was pressurised to 180 mmHg by injecting with air before imaging. All models could hold pressure with less than 10% loss over the eight minutes, recorded using a 0-15 psi gauge pressure sensor prior to imaging (Honeywell, Cat. No. 286-664). Experimental set-up and pressure results are presented in supplementary figure S3.

Radiographic data was processed using VGSTUDIO MAX (Volume Graphics, v2024.2) and exported in STL format for further processing in 3-matic software. Datum plane interactive cuts of the STLs allowed the inner lumen surface of each model to be extracted as a separate part. The resulting models were then aligned with the originally segmented artery to evaluate geometric accuracy, or with the un-pressurised model for analysis of distension. Part comparison analysis was conducted to produce a colour map of geometric differences. The Dice-Sørensen coefficient was calculated using the Segment Comparison module in 3D Slicer (v5.6.2) [30].

### 2.10 Endothelialisation of 3D aneurysm and arterial models

Barbed plugs were SLA printed in High Temp Resin (Cat. No: RS-C2-HTAM-02) and post-processed by washing in the FormWash for 6 minutes and in the FormCure for 120 minutes at 80°C. These were used to plug models and prevent spills and evaporation.

3D PDMS, Elastic50A and BioMed Elastic50A resin arterial models were rinsed with 70% (w/v) ethanol then distilled water and allowed to dry before oxygen plasma treatment at 70 W for three minutes. Models were sterilised by soaking in 70% w/v ethanol for ten minutes and exposed to germicidal ultraviolet light for 30 minutes. Samples were then washed twice with DPBS for five minutes and incubated at 37°C with 0.1% gelatine overnight.

The next day, 0.1% gelatine was aspirated, and HBECs cells seeded at 20,000 cells/cm^2^. Models were flipped after 60 minutes for even coverage. Media exchanges were performed every two days, with models fixed at day 7 in 4% paraformaldehyde for 30 minutes. Elastic50A resin models were then immersed in SBB for three hours and washed as described in section 2.5. 0.1% Triton X-100 was applied to permeabilise cells, followed by NucBlue^TM^ Fixed Cell ReadyProbes^TM^ Reagent (2 drops per mL in DPBS) and ActinGreen^TM^ 488 ReadyProbes^TM^ Reagent (2 drops per mL in DPBS) at room temperature for 30 and 60 minutes, respectively. Models were sectioned to facilitate imaging. Fluorescent images were acquired using a Nikon Eclipse Ti2 fluorescent microscope with analysis using ImageJ [28].

### 2.11 Statistical Analysis

Cell viability and IF studies were analysed for three biological replicates each with three technical replicates. Statistical analysis was performed using an unmatched one-way analysis of variance (ANOVA) with Welsh correction and pairwise comparisons using GraphPad Prism (v9.5.1). Geometric accuracy was analysed based on three independent aneurysm models with a two-way ANOVA conducted comparing aneurysm geometry and fabrication method. Results are expressed as a mean ± standard deviation, with p<0.05 accepted as statistically significant and indicated using a * (*p<0.05, **p<0.01). Figure 2 groups individual pairwise comparisons to improve clarity, where ** indicates all statistical comparisons within that group have a p-value less than 0.01.

**Figure 2:**
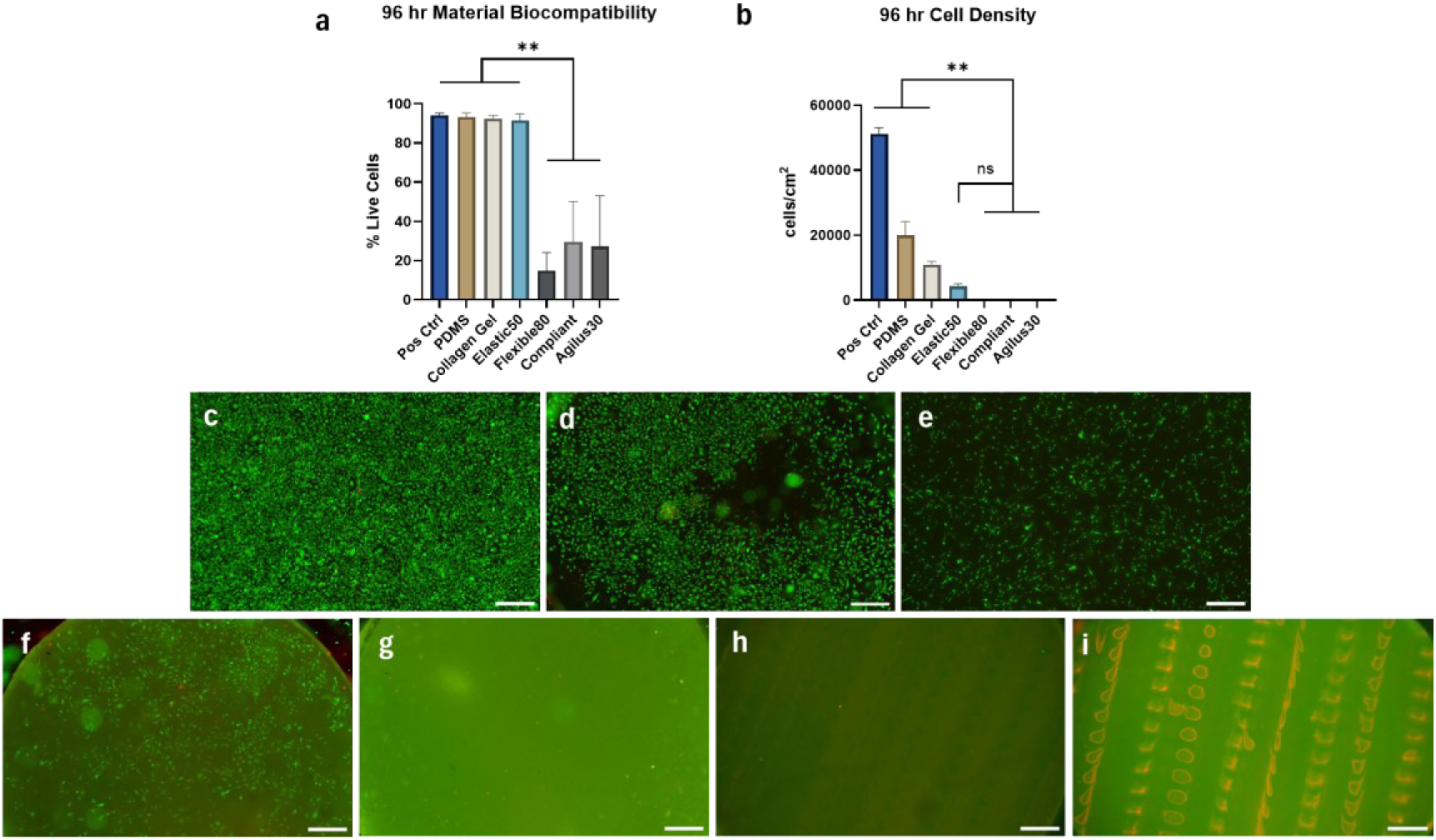
Additively manufactured material biocompatibility results at 96 hrs. (a) % live cells across materials. (b) Cell density on materials. (c) Positive control on tissue culture plastic. (d) PDMS. (e) Collagen Gel. (f) Elastic50A. (g) Flexible80A. (h) ‘Compliant’ blood vessel resin. (i) Agilus30. Human Brain Endothelial Cells (HBECs) cultured on materials were stained with Calcein-AM (green) and propidium iodide (red) and imaged at day four of culture. Scale bars represent 500 µm.

## 3 Results

### 3.1 HBEC viability

Cell viability was determined as the percentage of live cells stained with Calcein-AM to total combined cells stained with Calcein-AM and propidium iodide in a defined field of view. As shown in Figure 2, PDMS, collagen gel and the Elastic50A resin supported cell adhesion, achieving viabilities greater than 90%, with PDMS demonstrating the highest cell coverage at close to 20,000 cells/cm^2^. HBECs cultured on collagen gel were observed to adopt a spindle-like morphology compared to the polygonal spread-out morphology observed in the positive controls and PDMS cultures.

The Flexible 80A, ‘Compliant’ blood vessel resin, and Agilus30 resin substrates saw few cells attach, with any cells present appearing rounded and atypical in morphology. These materials were not investigated further due to their inability to support viable cell culture in this context.

### 3.2 Cell proliferation and coverage

PDMS, elastic 50A resin and collagen gel were further investigated for cell surface coverage and proliferation by IF staining for F-actin and Ki67. Ki67 expression was consistently above 50% in cultures on gelatine-coated tissue culture plastic, PDMS and elastic50A resin at 7 and 14 days of culture, with no significant difference between materials (Fig 3b). This consistently high expression observed on PDMS and tissue culture plastic despite the high cell density indicates HBECs are unlikely to undergo efficient contact inhibition, causing cells to continue to grow past confluence. Unfortunately, as can be seen in Figure 3A, there was significant non-specific Ki67 staining on collagen gel substrates, as evidenced in the isotype control stain. As a result, quantification of Ki67 expression was not feasible for this condition.

**Figure 3:**
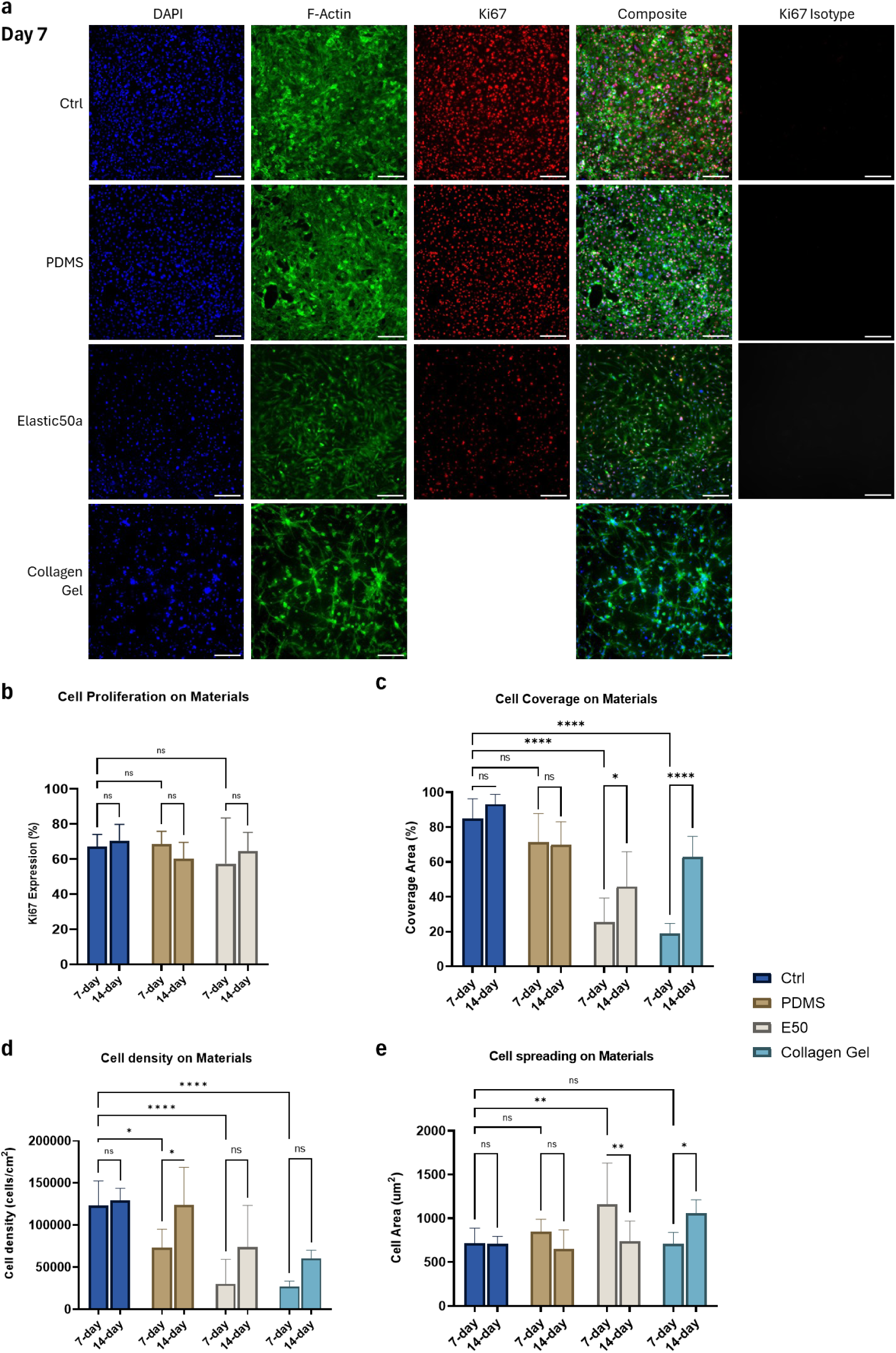
Morphology and proliferation of HBECs cultured for 7-days on tissue culture plastic (Ctrl), Polydimethylsiloxane (PDMS), elastic50A resin and collagen gel. Cells were stained for Ki67 (red), DAPI (blue) and F-actin (green) (a). Images analysed for cell growth (b), coverage area (c), cell density (d) and cell area (e). Due to non-specific binding of Ki67 to the collagen gel, these results could not be quantified. Scale bars represent 200 µm. Further images of and statistical comparisons between 14-day cultures can be found in Fig. S4.

High cell coverage (Fig. 3c) was consistently observed for PDMS at both 7– and 14-days of culture. A significant increase in surface coverage was observed on collagen gel and elastic50A resin at 14-days compared to 7 days, coinciding with an overall increase in cell density (Fig. 3d). Although, while cells tended to decrease in cell area as cell density increased (Fig. 3e), cells cultured on collagen gel were observed to expand. Images and statistical comparison of the 14-day cultures are presented in Supplementary Fig. S4.

### 3.3 Model Accuracy and Distensibility

Through µCT imaging, the accuracy of the fabrication process in producing the BA and MCA aneurysm models was evaluated and presented in Figure 4. The BA aneurysm could be reproduced using both the PDMS injection moulding and 3D printing techniques with an average DICE score of 90.1% and 94.5% respectively. However, the accuracy of the MCA aneurysm model fabricated in PDMS was significantly reduced (Average DICE 75.8%), compared to the basilar artery (P = 0.0142) and the 3D printed elastic50A version (average DICE 90.8%, P = 0.0187).

**Figure 4:**
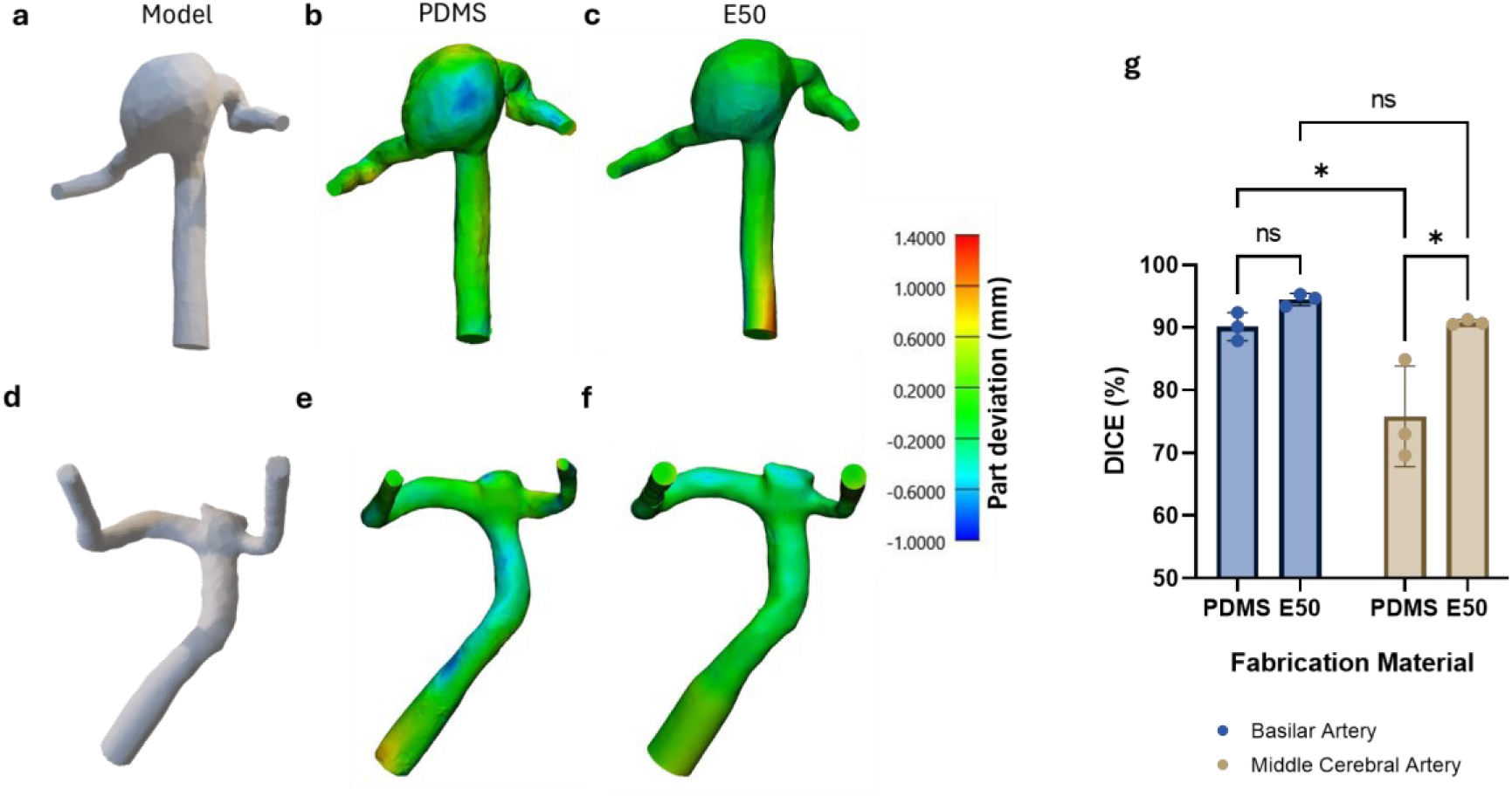
Manufacturing accuracy of aneurysm cell culture models. (a) Segmented basilar artery aneurysm. (b) Part deviation (mm) of the fabricated PDMS model compared to the original segmented basilar artery aneurysm. (c) Part deviation (mm) of the 3D printed Elastic50A resin model compared to the original segmented basilar artery aneurysm. (d) Segmented middle cerebral artery aneurysm. (e) Part deviation (mm) of the fabricated PDMS model compared to the original segmented middle cerebral artery aneurysm. (f) Part deviation (mm) of the 3D printed Elastic50A resin model compared to the original segmented middle cerebral artery aneurysm. (g) Two-way ANOVA analysis of Dice-SØrensen coefficient of fabricated aneurysm models.

The BA aneurysm model distension was then analysed when pressurised to approximately 180 mmHg using air. Figure 5a shows localised distention at the top of the aneurysm bulge in the PDMS model where the wall is thinnest, with no distension elsewhere due to the thick, block-like shape. Comparatively, the Elastic50A resin showed minimal expansion under pressure when printed with a 2 mm wall (Fig. 5b) while a 1 mm wall thickness saw some expansion across the aneurysm bulge (Fig. 5c). However, the arms of the model with a 1 mm wall were easily bent, causing significant deviation as shown in grey in the colour map of Figure 5c. The flexibility of these arms and inability to hold their shape could therefore significantly alter fluid dynamics within the aneurysm if used for *in vitro* perfusion studies.

**Figure 5:**
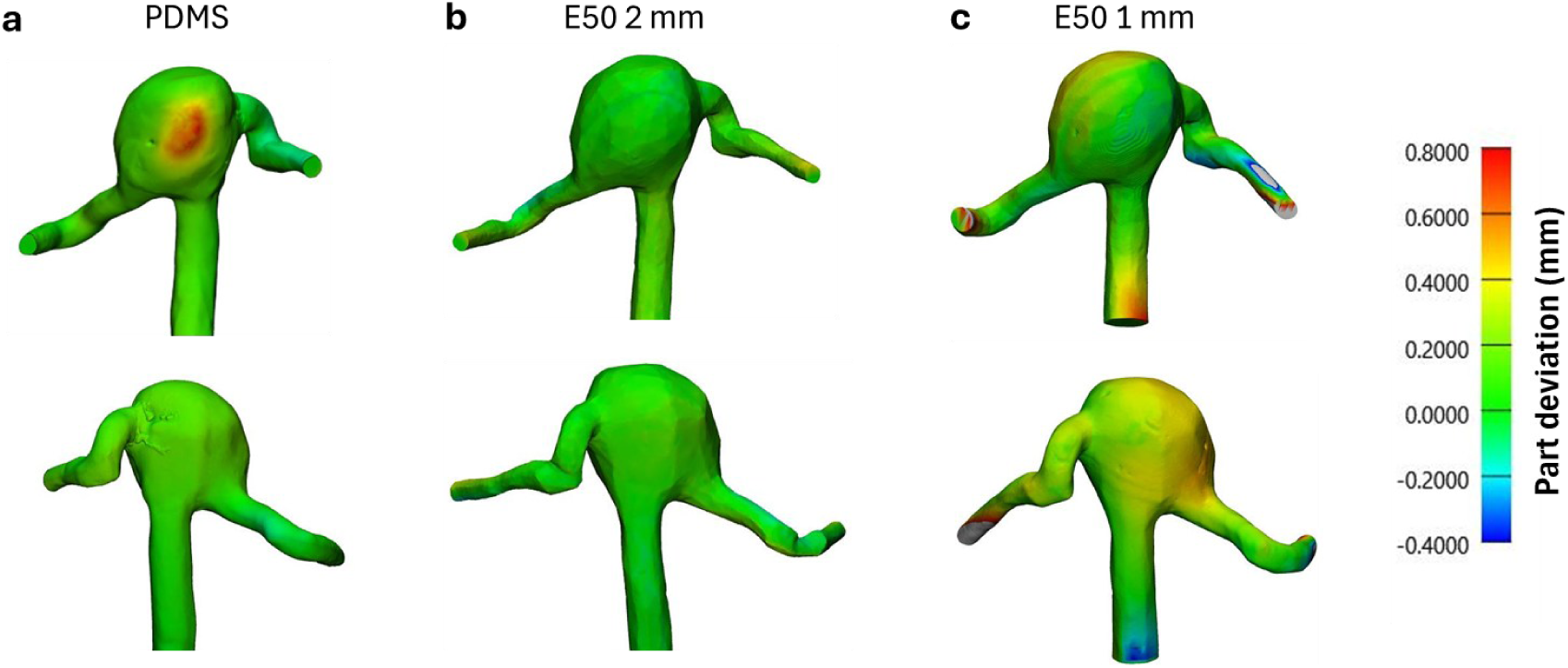
Distension of in vitro basilar artery (BA) aneurysm models. BA aneurysm models were pressurised to 180 mmHg. Part deviation heatmaps are presented for (a) chocolate injection-moulded PDMS models, (b) Elastic50A model with 2 mm wall thickness, and (c) Elastic50A model with 1 mm wall thickness.

### 3.4 Model Endothelialisation

HBECs were seeded into both the PDMS and elastic50A resin models to evaluate suitability of these materials as 3D *in vitro* models (Fig. 6 and Fig S5 respectively). IF staining for F-actin and DAPI saw complete coverage across the entirety of the basilar aneurysm PDMS model. No cells were seen in the elastic50A resin or BioMed elastic50A models after 7 days (Fig. S5), despite the BioMed elastic50A resin also demonstrating favourable results in preliminary trials in 2D (Supplementary Fig S6).

**Figure 6:**
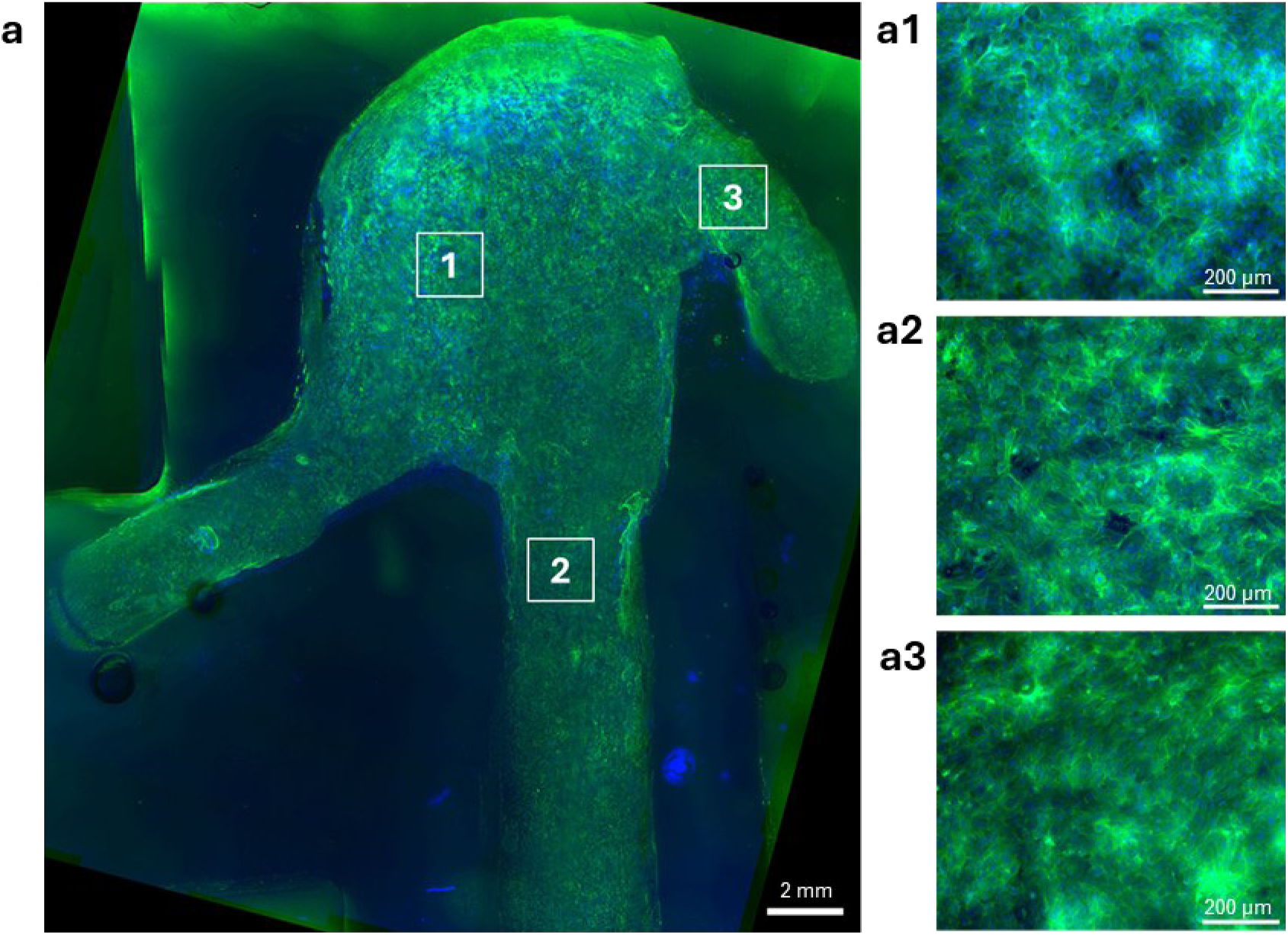
Human Brain Endothelial Cells (HBECs) cultured for 7-days in a 3D polydimethylsiloxane (PDMS) basilar artery aneurysm model. (a) stained for DAPI (blue) and F-actin (green).

## 4 Discussion

In this study, we evaluated two fabrication techniques for creating *in vitro* intracranial aneurysm models suitable for the cultivation of a human brain endothelial cell line. This included direct 3D printing, and the injection moulding of a chocolate sacrificial core which was embedded in PDMS and washed out. 3D printing benefits from its quick, easy and consistent fabrication process, while the injection moulded PDMS models included a time-consuming multi-step process subject to variation. Although, this novel chocolate-sacrificial core technique offers a non-toxic, support-free fabrication method enabling the use of a wide range of materials. Both PDMS and the 3D printed elastic50A resin supported viable cell attachment and expansion on material surfaces and could be fabricated into complex 3D patient-specific IA geometries. Despite the elastic50A resin proving viable in planar 2D cultures, only PDMS could support cell attachment and expansion in 3D IA models. This could be due to reduced gas permeability of the resin thereby reducing oxygen availability for cells. Comparatively, PDMS is well known for its oxygen permeability and thus widely used in tissue engineering applications [27]. These injection moulded PDMS IA models therefore present a promising platform for use in future research investigating cell-fluid interactions and IA pathophysiology.

The Formlabs flexible80A and Stratasys Agilus30 and ‘compliant’ blood vessel resins did not support viable cell attachment, consistent with previous reports [27, 31, 32]. Although, many commercial resins thought to be cytotoxic can be rendered biocompatible under specific post-processing techniques [27]. For example, Hart et al. (2020) showed that IPA sonication, UV-curing and autoclaving was sufficient to achieve 90% cell viability on the flexible80A resin, although the cell density retained on this material for analysis was not reported [33]. Further, Grigaleviciute et al. (2020) demonstrated prolonged exposure of the resin samples to IPA and methanol for over five days also significantly improved cell viability on this resin [34]. Therefore, specialised resin-specific post-processing methods could yield improved biocompatibility for the flexible80A, ‘compliant’ and Agilus30 resins than what was achieved herein [33, 34]. We found IPA washing, UV-curing, oxygen plasma treatment, ethanol sterilisation and gelatine coating of the elastic50A resin was a successful post-processing protocol to achieve high cell viability and growth in up to 14 days of culture. However, further investigation into the gas permeability of these resin materials is required to ensure suitability for use in closed cell culture models.

Collagen I, as a significant component of the cerebrovascular ECM was expected to offer optimal cell growth and viability in comparison to the synthetic PDMS and 3D-printed resins. However, cell growth was limited on collagen gel with morphologies ranging between rounded and spindle-like, compared to the polygonal spread-out morphology consistently observed on the positive controls and PDMS. Similar results were reported by Dye et al (2009), where human placental microvascular endothelial cells (HPMECs) adopted spindle-like morphologies and did not exhibit extensive cell-cell contact when grown on the surface of collagen I gels. Alternatively, the mechanical properties of a substrate is well known to influence the phenotype and response of cultured cells where Mason et al (2013) showed a decrease in cell area when embedded in collagen I gels of lower stiffness [35]. We observed an increase in cell area between 7– and 14-days on the collagen gel, which could be due to the increased cell-cell contact experienced, causing morphologies to become indistinguishable. Similar behaviours were observed in bovine aortic endothelial cells by Yeung et al. (2004) as cultures reached confluence [36]. This behaviour could also be a result of cell traction forces and ECM deposition contributing to gel stiffening over time [37, 38].

Further, Katt et al (2018) showed significant improvements in cell coverage on gels of high stiffness (up to 3.8 kPa) by increasing collagen gel concentrations [39, 40]. Comparatively, the type 1 atelocollagen gel used here offers a reported stiffness of approximately 200 Pa [41], while both PDMS and elastic50A yield a reported stiffness of approximately 1 and 3 MPa respectively [39–43]. Although, Katt et al (2018) demonstrated the increase in stiffness alone was not sufficient to form reliable monolayers on collagen gel, with gels requiring additional cross-linking with genipin or the supplementation of ROCK inhibitor or cAMP into culture medium to support monolayer formation [40]. While cell coverage can be improved on collagen gel, it was not considered further for 3D fabrication through the chocolate-sacrificial core technique due to the superior performance of PDMS.

Where model defects and curvature can influence fluid dynamics and cell behaviour, ensuring anatomical accuracy of fabricated IA models is essential for physiological relevance. While the 3D printed elastic50A resin IA models could be reproduced consistently and to a high accuracy (DICE > 90%) for both the BA and MCA models, the PDMS injection moulded technique was less reliable. Although the BA model could be fabricated with an average DICE score of 90.1% using this technique, the smaller MCA was less accurate at only 75.8% similarity. This is likely due to the smaller size of the MCA compared to the BA, making it prone to breakage and defects during the chocolate injection moulding process. Further, the shape of the MCA aneurysm bulge often caused air pockets to form during injection moulding, impeding proper formation of the aneurysm. 3D printed IA models used for surgical simulations have previously been analysed for printing accuracy, reporting DICE scores between 84% and 97% [44–46]. These studies also demonstrate a reduction in DICE score with reduced aneurysm volume. Both Haruma et al. (2022) and Shibata et al. (2017) propose a DICE score above 70% indicates acceptable printing performance, of which both PDMS injection moulded IA models exceed.

While the effects of mechanical stretching on endothelial cells have been evaluated in specialised devices [47, 48], the effects of cyclic stretching on cell behaviour in *in vitro* IA models has not yet been investigated. This is important where aneurysms can present with a thin and/or thick-walled phenotype which can influence the degree of cyclic stretching [49]. For example, decreased arterial distensibility and increased intima-media thickness has been observed in patients with ruptured IAs [50]. We therefore demonstrate the ability to tune model distension behaviour through varying IA model wall thickness. Where a thin wall was designed at the BA aneurysm bulge of the PDMS model, localised expansion of up to 0.8 mm was observed when pressurised to 180 mmHg.

Although 3D printed resin IA models were found to be unsuitable for this application using the current processing techniques, we present an alternative fabrication method for developing PDMS IA models with validated anatomical accuracy. We demonstrate that modulating wall thickness of these models could enable the evaluation of localised effects of cyclic strain combined with haemodynamic stress.

This study on *in vitro* IA models has several limitations. First, the sacrificial chocolate core can be delicate and prone to breakage, particularly for smaller geometries. For example, the asymptomatic MCA geometry was unable to be fabricated due to the small diameter at the bifurcation (approximately 1.8 mm) frequently breaking. The time-consuming multi-step process also increases difficulty and risk of defects. Prolonged manual handling can cause slight melting and deformation of the models over time due to the low melting temperature. Although, by heating a spatula slightly, small defects can be smoothed easily without significantly altering the shape. Alternative sacrificial core materials were explored, for instance using a 3D printed polyvinyl alcohol (PVA) core which was difficult to process and remove supports without breaking, while an injection moulded water-soluble wax core also proved fragile.

Secondly, as all 3D printed resins used are photopolymers and typically cured under UV (405 nm) light, they preclude IF analysis under the same wavelength, such as when staining for DAPI (360 – 460 nm). Sudan Black B was thus used to quench the autofluorescence of elastic50A resin in IF analyses. While this revealed cell nuclei, it reduced signal of F-actin under the FITC wavelength filter (488 nm), requiring additional image post-processing.

Third, the basilar artery models were a simplification of the true geometry of the basilar artery, as side branching pontine arteries were removed. This is a common practice in CFD modelling where smaller perforating arteries are seldom included in simulations, or where boundary conditions are unknown and difficult to acquire [51]. Therefore, care should be taken when evaluating cell response to fluid flow through these geometries due to the currently unknown influence of these side branches on fluid flow patterns.

While simple, idealised IA models have previously been developed in hydrogels, no complex patient-specific IA geometries have been created [16]. This could be achieved using the sacrificial chocolate core method described here, which benefits from being non-toxic with a relatively low melting temperature. However, due to the superior results of PDMS compared to collagen gel, we did not investigate this further. Alternative hydrogels such as GelMA or fibrin-based gels could be suitable alternatives for embedding a sacrificial chocolate core to create patient-specific IA models.

Furthermore, evaluating endothelial cell behaviour has been the focus of this study as it plays a key role in the formation, growth and rupture of an IA. However, IA pathophysiology is complex and influenced by a range of factors including smooth muscle cells and fibroblasts, and changes in the extracellular matrix and inner elastic lamina. Future *in vitro* IA models can extend on this study to incorporate the structural and cellular complexities of the IA to accurately represent a human IA.

Finally, BA models were pressurised to hypertensive conditions (maximum 180 mmHg) as an exploratory study to exemplify maximum mechanical strain achievable [52]. Under normotensive conditions, these models would be exposed to maximum pressures of approximately 113 mmHg at a blood flow rate of around 145 mL/min typically seen in the basilar artery [52, 53].

## 5 Conclusion

Biofabricated IA cell culture models are promising platforms for investigating cell-fluid interactions to improve understanding of IA pathophysiology. By evaluating several fabrication techniques and materials, we found that PDMS remains the superior material for fabricating 3D patient-specific IA geometries suitable for cell culture. These models can be fabricated by moulding PDMS around a patient-specific sacrificial core made of tempered chocolate, achieving shape-fidelities above 90%. This novel chocolate-sacrificial core offers a non-toxic, support-free fabrication method suitable for 3D cultures. While the elastic50A resin was unsuitable for enclosed 3D applications such as arterial models, it proved viable for open planar material surface cultures despite previous reports of cytotoxicity. We further demonstrated the ability to create localised distension in these models by modulating wall thickness, making it useful for studying IA pathophysiology in the context of thin– and thick-walled aneurysms.

Future studies could see these patient-specific PDMS IA models cultured with human vascular endothelial cells and subject to physiological fluid flow. Cell response to haemodynamic stress and cyclic strain in the context of intracranial aneurysms can then be investigated.

## 6 Acknowledgements

The authors would like to thank the Royal Brisbane and Women’s Hospital (RBWH) Foundation and the RBWH Department of Medical Imaging for their support, including RBWH Foundation grants in 2020 and 2021. The authors would also like to thank Metro North Health who funded materials and some of the researchers through the CranioFacial program of the Herston Biofabrication Institute. This work was supported by the Australian Research Council (DE220100757) and Advance Queensland (AQIRF1312018) fellowships. This work also used the Queensland node of the NCRIS-enabled Australian National Fabrication Facility (ANFF).

The authors would like to thank Roozbeh Fakhr, Liam Georgeson, Kaecee Fitzgerald and Dr David Forrestal from the Herston Biofabrication Institute for their assistance in 3D modelling, printing and processing. Dr Ekaterina Strounina from the University of Queensland Centre for Advanced Imaging, part of the National Collaborative Research Infrastructure Strategy (NCRIS) and the National Imaging Facility (NIF), for her assistance with µCT imaging and processing. Professor Justin Cooper-White and Ms Taryn Smith from the University of Queensland for the use of their pressure transducer and for aiding pressure monitoring studies.

## SUPPLEMENTARY MATERIAL

**Figure S1:**
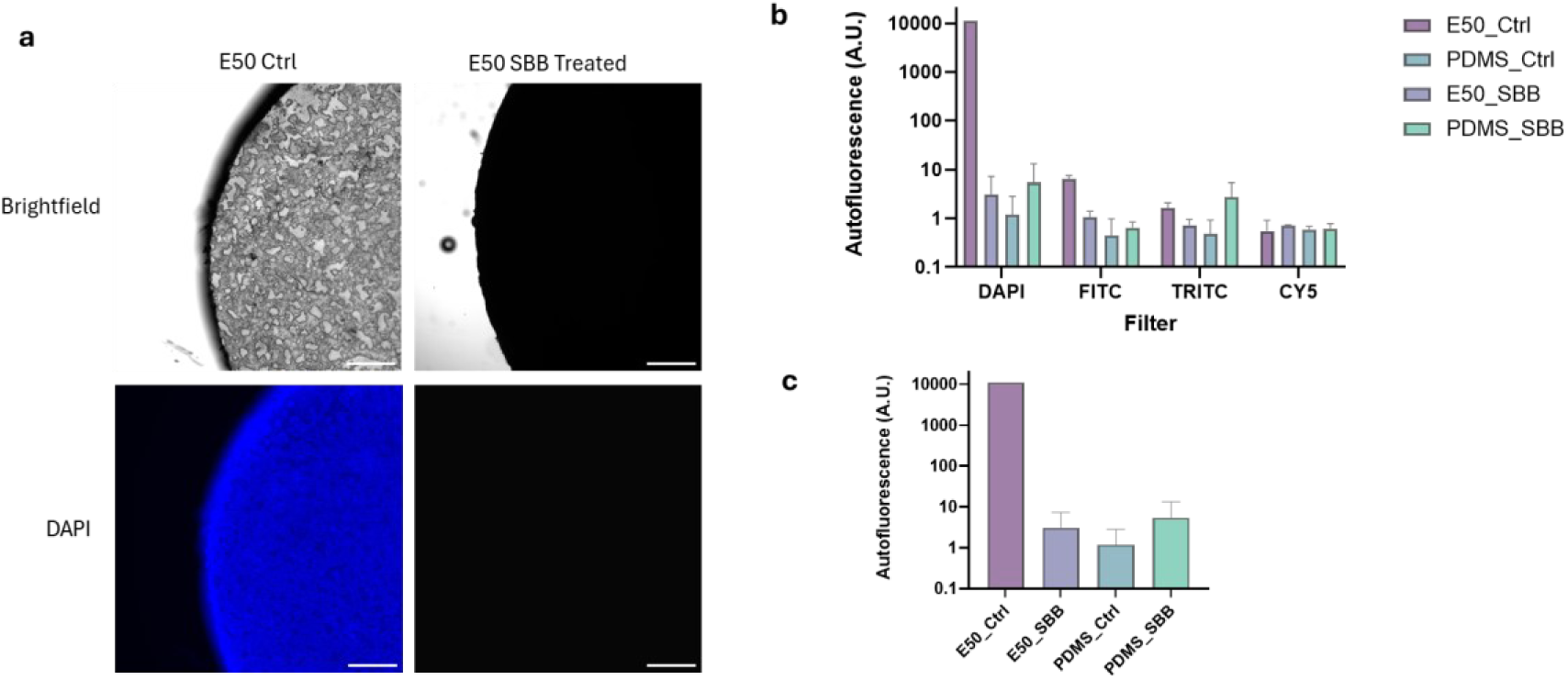
Sudan Black B reduces Elastic50A resin autofluorescence. (a) Comparison of Elastic50A (E50) resin under brightfield and DAPI fluorescent filters when untreated and treated with Sudan Black B SBB. (b) Evaluation of autofluorescence through pixel intensity of E50 and PDMS materials under DAPI, FITC, TRITC and CY5 fluorescent filters. (c) Autofluorescence of E50 and PDMS materials under the DAPI fluorescent filter. Scale bar represents 500 µm.

**Figure S2:**
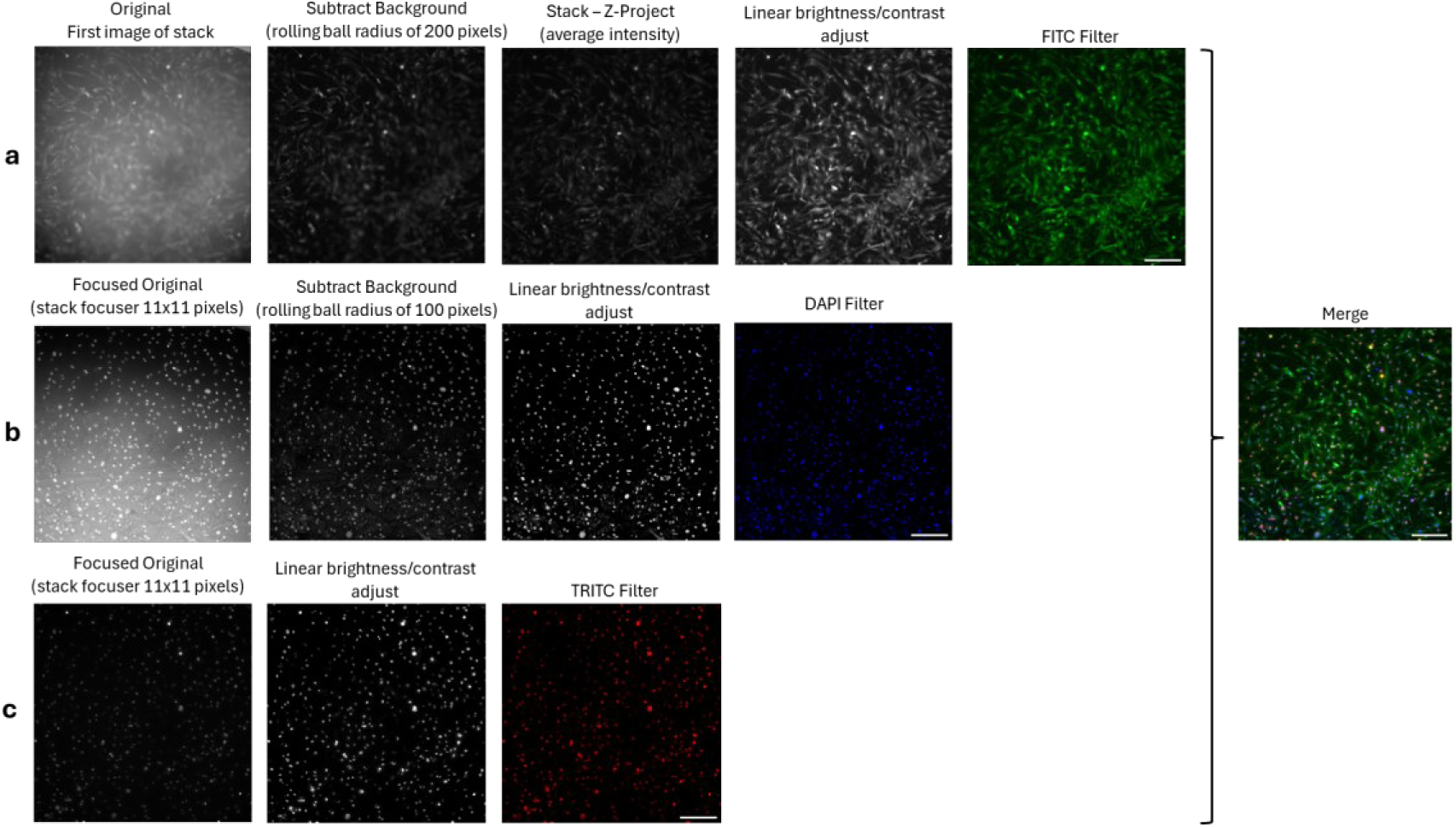
Image processing of elastic50A resin samples treated with Sudan Black B using ImageJ. (a) Processing sequence for images obtained in the FITC fluorescent filter of cells stained for F-actin. (b) Processing sequence for images obtained in the DAPI fluorescent filter of cells stained for the cell nucleus. (c) Processing sequence for images obtained in the TRITC fluorescent filter of cells stained for Ki67. Scale bar represents 200 µm.

**Figure S3:**
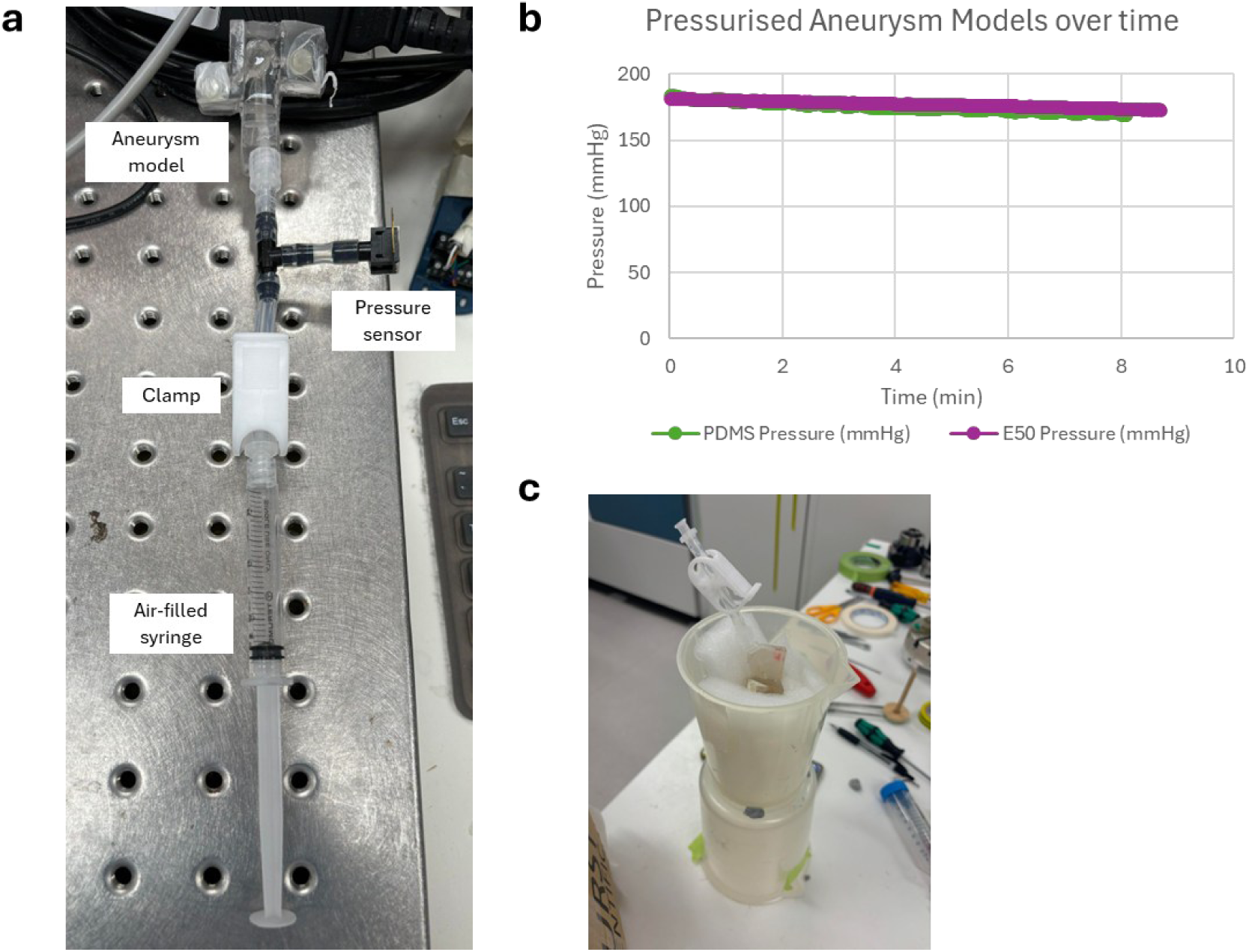
Aneurysm model air pressurisation set-up and measurement. (a) PDMS basilar artery aneurysm model gas pressurisation setup connected to a 0-15 psi gauge pressure sensor (Honeywell, Cat. No. 286-664). (b) Recorded pressure overtime within PDMS and elastic50A resin basilar artery aneurysm models. Recorded before µCT imaging. (c) Aneurysm model placed in tub with foam to prevent movement during µCT imaging. Models were pressurised with an air-filled syringe, then clamped and the syringe removed before imaging.

**Figure S4:**
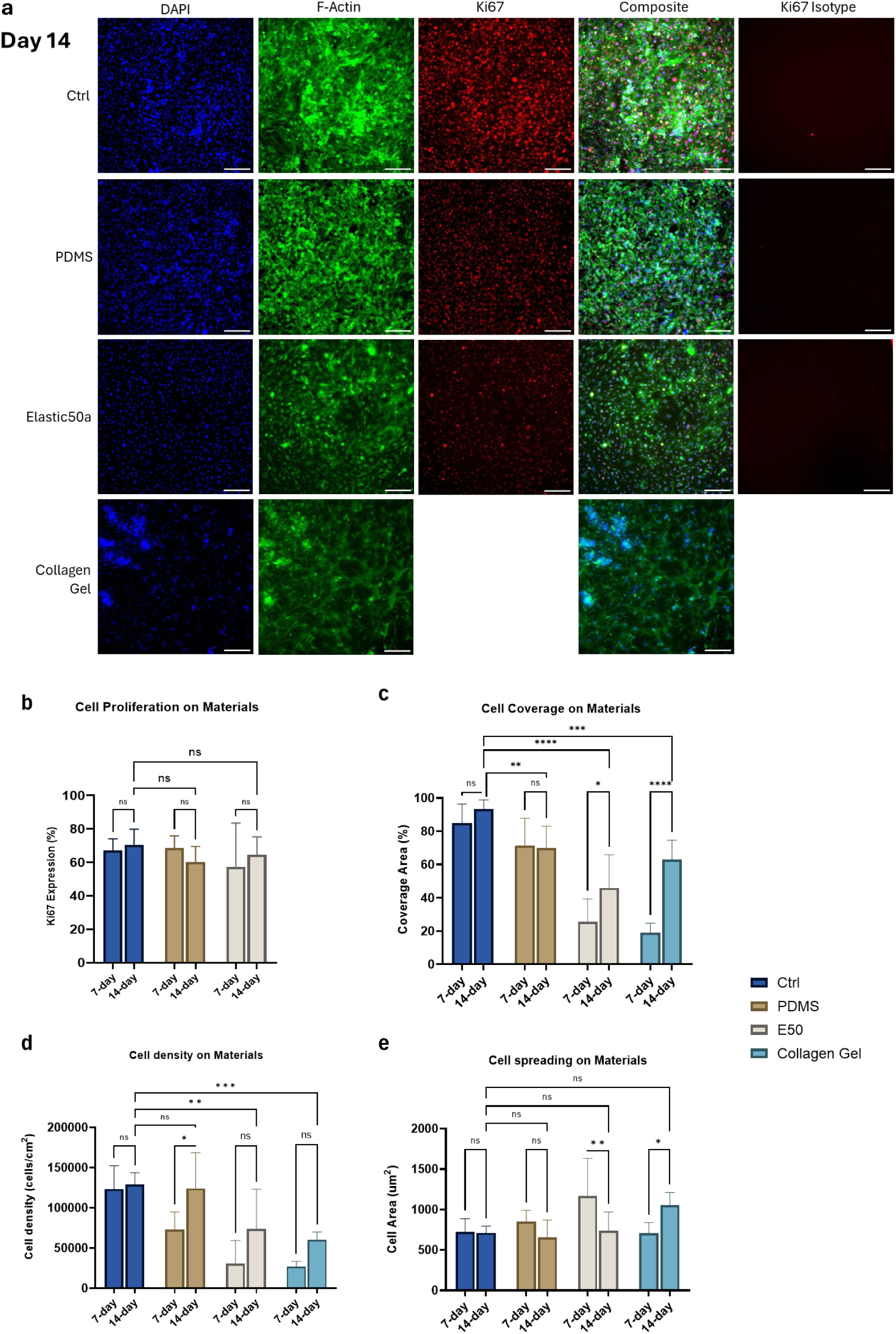
Additional statistical comparisons of HBECs cultured for 14-days on tissue culture plastic (Ctrl), Polydimethylsiloxane (PDMS), elastic50A resin and collagen gel. Cells were stained for Ki67 (red), DAPI (blue) and F-actin (green) (a). Images analysed for cell growth (b), coverage area (c), cell density (d) and cell area (e). Due to non-specific binding of Ki67 to the collagen gel, these results could not be quantified. Scale bars represent 200 µm.

**Figure S5:**
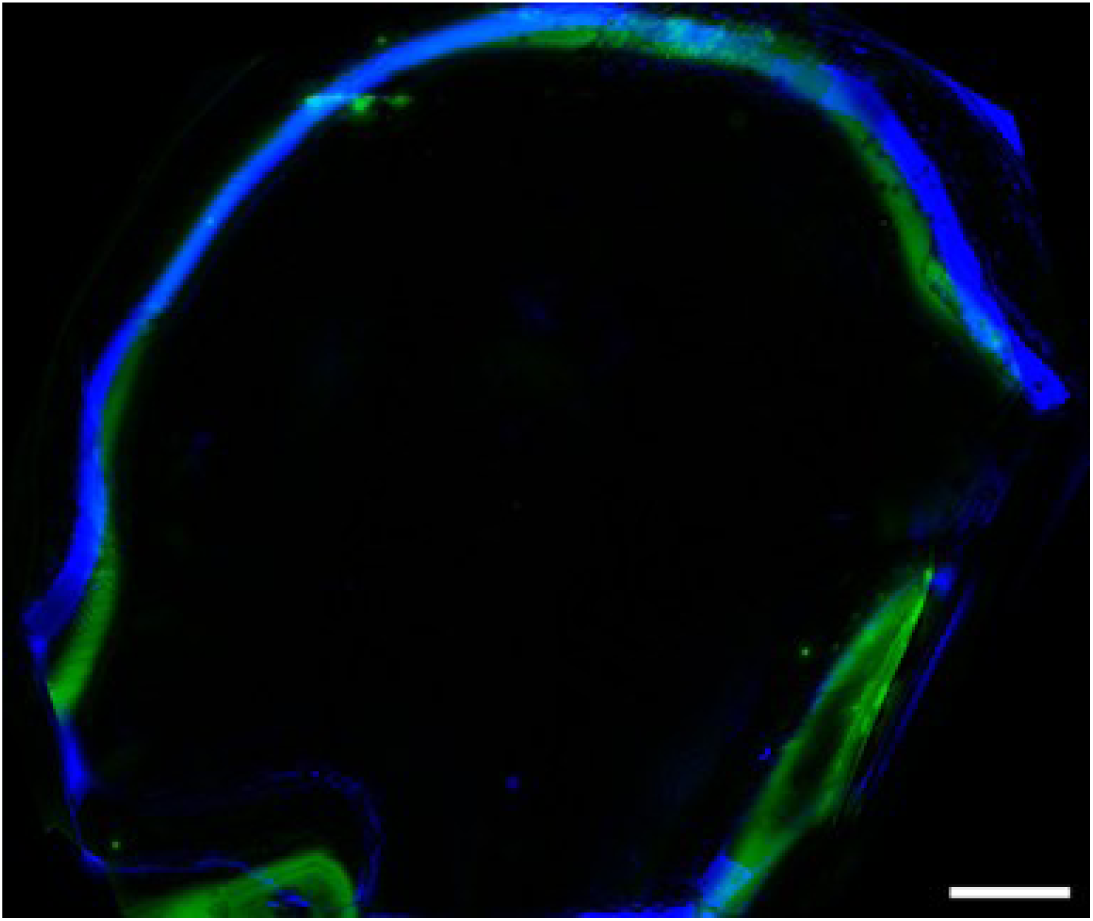
Immunofluorescence of Human Brain Endothelial Cells (HBECs) cultured for 7 days in a 3D basilar artery model made from Elastic50A resin. Stained for DAPI (blue) and F-actin (green). Sudan black B (SBB) was applied to the models prior to sectioning to quench surface autofluorescence. While SBB quenched autofluorescence at the surface of the elastic50A aneurysm lumen, the walls of the sectioned model not exposed to SBB continued to fluoresce. Scale bar represents 2 mm.

**Figure S6:**
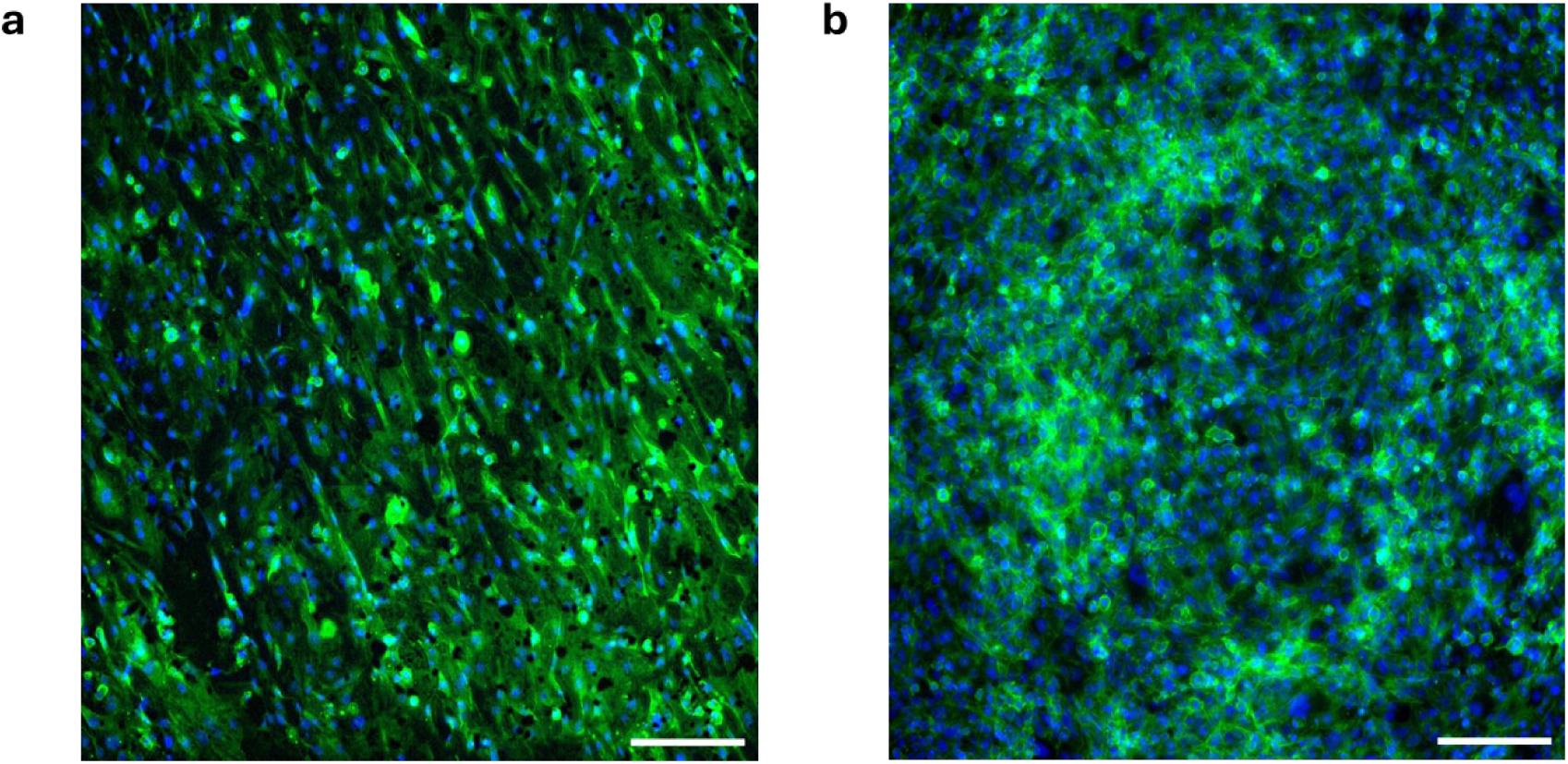
Preliminary analysis of the suitability of BioMed E50 resin for the culture of Human Brain Endothelial Cells (HBECs) over 7 days. (a) BioMed E50 resin cultured with HBECs for 7 days. Material was printed on a 45° angle producing layer lines impacting cell coverage and alignment. Image processed as per processing steps outlined in Figure S2. (b) Tissue culture plastic control. Scale bar represents 200 µm.

